# Piperacillin-Tazobactam Resistance Mechanisms in *Escherichia coli* and Identification of a CTX-M-255 β-Lactamase Selectively Conferring Resistance to Penicillin/β-Lactamase Inhibitor Combinations

**DOI:** 10.1101/2022.09.21.508968

**Authors:** Minna Rud Andreasen, Katrine Hartung Hansen, Martin Schou Pedersen, Sarah Mollerup, Lotte Jelsbak, Kristian Schønning

## Abstract

Piperacillin/tazobactam (TZP) is a widely used penicillin/β-lactamase inhibitor combination with broad antimicrobial activity. Recently, *Escherichia coli* strains resistant to TZP but susceptible to third generation cephalosporins (TZP-R/3GC-S isolates) have been increasingly identified. Here, we investigated resistance mechanisms underlying the TZP-R/3GC-S phenotype in clinical *E. coli* isolates.

A total of 29 TZP-R/3GC-S *E. coli* isolates were retrieved from urinary cultures and subjected to whole genome sequencing. Resistance to TZP was confirmed by minimum inhibitory concentration determination. β-lactamase activity in the presence and absence of tazobactam was determined to identify hyperproduction of β-lactamase and assess susceptibility to tazobactam inhibition. A previously unrecognized β-lactamase was identified and cloned to determine its resistance profile.

Four different resistance mechanisms underlying the TZP-R/3-GC phenotype were identified: 1) In 18 out of 29 isolates (62%) β-lactamase production was increased and in 16 of these either strong alternative promoters or increased gene copy numbers of *bla*_TEM-1_ or *bla*_SHV-1_ were identified, 2) seven isolates (24%) produced *bla*_OXA-1_, 3) three isolates (10%) produced inhibitor-resistant TEM-β-lactamases, and 4) a single isolate (3%) harboured a *bla*_CTX-M_ gene as the only β-lactamase present. This β-lactamase, CTX-M-255, only differs from CTX-M-27 by a G239S amino acid substitution. In contrast to CTX-M-27, CTX-M-255 conferred resistance to penicillin/β-lactamase inhibitor combinations but remained susceptible to cephalosporins.

In conclusion, hyperproduction of *bla*_TEM_ was the most prevalent mechanism of TZP-resistance underlying the TZP-R/3GC-S phenotype followed by production of *bla*_OXA-1_ and inhibitor-resistant TEM-β-lactamases. Furthermore, we identified a previously unrecognized CTX-M-β-lactamase, CTX-M-255 that was resistant to β-lactamase inhibitors.

## Introduction

Piperacillin/tazobactam (TZP) is a penicillin/β-lactamase inhibitor combination widely used in hospitals for treatment of severe infections. In Switzerland, Germany, Canada, and Denmark, TZP accounted for 7.8% (2017), 12.1% (2016), 12.3% (2018), and 11.6% (2020) of antimicrobial consumption in hospitals, respectively (1–4). The resistance rate of TZP has thus far remained relatively low in Denmark with a TZP resistance of 4.3% of all hospital urine *Escherichia coli* isolates in 2020 (4). However, given its wide usage and thus risk of resistance development, an understanding of resistance mechanisms to TZP is important.

Acquired TZP resistance is often accompanied by resistance to third generation cephalosporins and may be caused by acquisition of β-lactamases such as AmpC enzymes, extended spectrum β-lactamases, or carbapenemases (5–7). However, TZP resistant isolates of *E. coli* and *Klebsiella pneumoniae* that remain fully susceptible to third generation cephalosporins (TZP-R/3GC-S) have also been reported (8–10). Prevalence of TZP-R/3GC-S isolates of 4.2% and 5.8% in *E*.*coli* and *K. pneumoniae* blood stream infections have been reported from hospitals in New York (USA) and Liverpool (UK), respectively (8, 10) and TZP-R/3GC-S isolates constituted 50% of all TZP resistant isolates (10).

We have previously investigated TZP resistance mechanisms in a set of six TZP-R/3GS-S *E. coli* isolates, in which *bla*_TEM_ was identified as the only acquired β-lactamase (11). In five of these isolates, hyperproduction of β-lactamase was observed as a cause of TZP-resistance and associated with either *bla*_TEM-1_ gene amplifications or strong *bla*_TEM-1_ promoter variants: *Pa*/*Pb, P4*, or *P5* (12). Both gene amplification and strong promoters of *bla*_TEM-1_ caused increased production of TEM-1 (11). Indeed, the degree of hyperproduction of *bla*_TEM-1_ by these isolates correlated with the concentration of tazobactam required to restore piperacillin susceptibility (13). The last isolate co-produced *bla*_TEM-1_ and *bla*_TEM-35_ the latter being an inhibitor resistant TEM (IRT) enzyme not efficiently inhibited by tazobactam and a well described cause of penicillin/β-lactamase inhibitor resistance (14–16). Similar to *bla*_TEM-1_ hyperproduction, TZP resistance can be caused by hyperproduction of *bla*_SHV-1_ due to increased copy number (9) or a stronger promoter (17).

*bla*_OXA-1_ is a class D enzyme and is not inhibited by most β-lactamase inhibitors, e.g. clavulanic acid and tazobactam (18). In a Spanish survey of *E. coli* isolates resistant to amoxicillin/clavulanate, expression of *bla*_OXA-1_ was attributed as the causative resistance mechanism in 21.0% of the isolates (16). Furthermore, in a study of ESBL-carrying *E. coli* co-carriage of *bla*_OXA-1_ correlated with resistance to penicillin/β-lactamase inhibitor combinations including TZP (19).

In the present study, we investigated 29 TZP-R/3GC-S *E. coli* isolates to estimate the relative contributions of 1) hyperproduction of *bla*_TEM-1/SHV-1_, 2) IRT enzymes, and 3) carriage of *bla*_OXA-1_ to the observed phenotype. Furthermore, in an isolate in which none of the above resistance mechanisms was present, a previously undescribed β-lactamase, CTX-M-255, was identified, which was resistant to inhibition by β-lactamase inhibitors.

## Materials and methods

### Study isolates

Clinical *E. coli* isolates from urinary tract infections resistant to TZP and susceptible to cefpodoxime, cefoxitin, and temocillin were isolated from urine samples referred to analysis at Department of Clinical Microbiology, Hvidovre Hospital, from December 2017 to May 2018. Susceptibility to cefoxitin and temocillin was chosen as inclusion criterion to exclude TZP resistant isolates producing AmpC β-lactamases and OXA-48-like β-lactamases, respectively. Only one isolate was included if multiple isolates were obtained from the same patient. Isolate EC120 was previously sequenced with Oxford Nanopore Technology (ONT) and characterized as part of a previously published work, where TZP-R/3GC-S *E. coli* isolates were screened for tandem repeats of *bla*_TEM_ by PCR and EC120 turned up positive (11). Isolate EC101, functioned as TZP susceptible control producing TEM-1 in low levels with a weak *P3* promoter and a *bla*_TEM-1_ copy number of 1.2 with the resistance phenotype: susceptible to TZP, amoxicillin/clavulanate, mecillinam, cefpodoxime, cefoxitin and temocillin but resistant to ampicillin.

### Antimicrobial susceptibility testing

Susceptibility to amoxicillin/clavulanate, mecillinam (amdinocillin), cefpodoxime, cefoxitin and temocillin was determined using disc diffusion and EUCAST methodology and interpreted using EUCAST clinical breakpoints table version 12.0. Further, TZP minimum inhibitory concentration (MIC) was determined by the broth microdilution method in accordance with EUCAST with piperacillin concentrations from 1 to 1024 mg/L supplied with 4 mg/L tazobactam. In addition, MIC susceptibility determination of *E. coli* TOP10/pZS3(*) transformant strains was performed with ampicillin, amoxicillin, and piperacillin alone and combined with β-lactamase inhibitors sulbactam, tazobactam, and avibactam (all fixed concentrations of 4 mg/L) and clavulanic acid (fixed concentration of 2 mg/L). Product information of all antibiotics is available in Table S1. All MICs were determined in duplicates.

### Whole genome sequencing

Whole genome sequencing of all 29 isolates was done as previously described and Illumina reads *de novo* assembled into contigs by SPAdes v. 3.11.1 (11). Multi Locus Sequence Typing (MLST) was performed by the open service https://cge.food.dtu.dk/services/MLST/ (20). Core genome MLST (cgMLST) was analyzed using the EnteroBase (http://enterobase.warwick.ac.uk) Escherichia/Shigella cgMLST v1 scheme with 2513 target genes implemented for *E*. *coli* in Ridom Seqsphere+ (Ridom GmbH, Munster, Germany, http://www.ridom.de/seqsphere/) and a distance matrix showing pairwise allele distances between the isolates was created. Acquired resistance genes associated with resistance to β-lactam antibiotics were found by ResFinder version 4.0 (21). Copy numbers of the resistance genes *bla*_TEM_, *bla*_SHV-1_, and *bla*_OXA-1_ were estimated by calculating the depth of coverage of these genes relative to the mean coverage depth of the seven *E. coli* MLST genes after mapping reads to these with Bowtie2. Promoter analysis of *bla*_TEM_ and *bla*_SHV-1_ was done manually in Geneious Prime v. 2021.2.2 following the proposed promoter nomenclature by Lartigue et al. (12). If examination of a β-lactamase-containing contig showed less than 300 base pairs upstream of the start codon, Illumina raw reads were examined and mapped to the reference genes to search for single nucleotide polymorphisms (SNPs) in the raw reads. A SNP within the promoter region or resistance gene itself covered by only a subset of reads is suggestive of multiple loci of the resistance gene in question. Such isolates were additionally sequenced with ONT as previously described (11). ONT long reads were used to unambiguously link individual β-lactamase loci to their respective promoter. Sequence data from this study has been deposited at NCBI BioProject PRJNA855633.

### β-lactamase activity determination

Cells from overnight cultures of the 29 TZP resistant isolates and TZP susceptible control isolate, EC101, were grown in Luria-Bertani broth at 37°C with shaking (150 rpm) were harvested and resuspended in 10 µL PBS/µg cells. Cells were lysed by sonication at 3×10 seconds at 35 kHz output and kept on ice for 5 min. Cellular debris was removed by centrifugation and the supernatant collected (lysate). β-lactamase activity determination in duplicates of untreated and 5 mg/L tazobactam treated (90 minutes at room temperature) lysates were performed using the Amplite™ Colorimetric β-lactamase Activity Assay Kit (AAT Bioquest, CA, USA) according to manufacturer’s protocol. β-lactamase activities were normalized relative to total protein concentration measured using Pierce Coomassie Plus Assay Kit (Bio-Rad, CA, USA) to obtain the enzyme activity in units pr. total protein (mU/mg).

### Construction of *E. coli* TOP10/pZS3(*) transformants

Two low-copy plasmids pZS3-CTX-M-15 and pZS3-CTX-M-178 (CTX-M-15_G239S) were generously supplied by Rosenkilde et al. (22). These plasmids express CTX-M-15 and CTX-M-178 under the control of a *bla*_TEM_ *P3* promoter and have a *bla*_TEM_ terminator downstream of the β-lactamases. To compare and characterize the resistance of the previously unrecognized CTX-M-255 we constructed pZS3*-CTX-M-255 and pZS3*-CTX-M-27 transformants in the same low-copy plasmid with the same terminator but with a native CTX-M promoter commonly associated with CTX-M-27 and identical to the promoter controlling CTX-M-255 in the parental isolate, WGS1364. These two plasmids have an asterisk in pZS3* to mark that the promoter region differ from the *P3* promoter of pZS3 plasmids. pZS3* have a 210 bp promoter region with an IS*Ecp1* element 42 bp upstream of the start codon of CTX-M. For this construction, the backbone of pZS3 ancestor (also supplied by Rosenkilde et. al.) was amplified and linearized by PCR with primer pair Fw-pZS3 backbone and Rv-pZS3 backbone (Table S2), thus consisting of a 120 bp fragment upstream of the *P3* promoter (to ensure the same length of the plasmids), plasmid core, and the *bla*_TEM_ terminator. The PCR product was treated with Dnp1 enzyme (ThermoFisher, MA, USA) to remove template plasmid and subsequently purified (QIAquick PCR Purification Kit, Qiagen, Germany). Two gene fragments were synthesized (Twist Bioscience, CA, USA) and consisted of 210 bp promoter sequence and either *bla*_CTX-M-27_ or *bla*_CTX-M-255_ and 29-30 bp homolog to the amplified pZS3 backbone at each end of the fragment. GeneArt TM Gibson Assembly HiFi Master Mix (ThermoFischer #A46628) was used for cloning the two gene fragments into pZS3 thereby contructing pZS3* by homologous overlapping ends according to the manufacturer’s protocol. All two pZS3 and two pZS3* plasmids were transferred to and maintained in *E. coli* TOP10 (ThermoFisher) with 50 mg/l kanamycin. The structure of the pZS3-CTX-M-15/178 and pZS3*-CTX-M-27/255 plasmids are visualized in Figure S1 and the promoter sequences are available in Table S2.

### Prediction of the 3D protein structure of CTX-M-255 and CTX-M-178

The protein structure of the wildtype CTX-M-27 protein was previously resolved by X-ray crystallography accessible at Protein Data Bank with accession ID: 1YLP. To predict and model the structure of the novel CTX-M-27-like β-lactamase, CTX-M-255, the online protein prediction server, Robetta (https://robetta.bakerlab.org/) was utilized where 1YLP served as template for comparative modeling of CTX-M-255. Likewise, CTX-M-178 (CTX-M-15_G239S) was modelled using CTX-M-15 (PDB ID: 4HBU) as template.

## Results

This study included 29 TZP-R/3GC-S *E. coli* isolates that were subjected to whole genome sequencing. From the sequences obtained MLST types were assigned. Six isolates were ST73, and six isolates were ST131. The remaining isolates represented different MLST types. Allele distance of *E. coli* cgMLST genes revealed that the isolates were genetically unrelated except EC123 and EC140 for which an allele distance <20 was observed (Table S3). EC123 and EC140 were isolated from different patients at separate hospitals 41 days apart. The two isolates were thus not epidemiologically related, and both were kept in this study.

The sequence data obtained was analysed for the presence of β-lactamase encoding genes. In 19 of 29 isolates, only variants of *bla*_TEM_ were identified. In five of these isolates, the presence of multiple alleles of *bla*_TEM_ was suspected from the assembly and subsequent read mapping analysis. These isolates were subjected to ONT sequencing, which confirmed the presence of multiple alleles and was used to link individual TEM-variants to their promoter.

In 16 of 29 isolates hyperproduction of TEM-1 (n=14) or SHV-1 (n=2) were identified as the likely causes of TZP resistance. Eight of 16 isolates had strong promoters associated with *bla*_TEM-1_ genes as likely cause of the observed TZP resistance. In six of these isolates, *bla*_TEM-1_ was associated with a strong *Pa*/*Pb* promoter, and in two isolates, *bla*_TEM-1_ was associated with hybrid promoters created by the insertion of IS*Apl1* or IS*26* elements within the *P3* promoter contributing alternative strong promoters. However, the isolate with an IS*26*/*P3* hybrid promoter had a *bla*_TEM-1_ copy number of 42.7 indicating that both a strong promoter and an increased copy number likely contribute to hyperproduction of *bla*_TEM-1_ and thus resistance to TZP. Six of the 19 isolates harboured *bla*_TEM-1_ associated with a *P3* promoter as the only β-lactamase present in high copy numbers ranging from 13.4 to 54.6 relative to MLST genes, constituting a likely explanation for the observed TZP resistance. Two of the 29 isolates harboured *bla*_SHV-1_ in increased copy numbers of 39.5 and 49.1, likely explaining the observed TZP resistance. In addition to TZP resistance all 16 isolates hyperproducing TEM-1 or SHV-1 were also resistant to mecillinam.

Three of 29 (10%) isolates contained IRT β-lactamases. EC133 contained IRT *bla*_TEM-30_ with a strong *Pa/Pb* promoter as the only β-lactamase. As previously reported, EC120 contained five copies of IRT *bla*_TEM-35_ with a *P4* promoter and a single copy of *bla*_TEM-1B_ with a *P3* promoter at different chromosomal loci (11). The last IRT isolate, EC126, contained three *bla*_TEM_ alleles; an allele containing IRT *bla*_TEM-32_ controlled by a *Pa*/*Pb* promoter, a *bla*_TEM-1B_ allele with a *Pa/Pb* promoter, and finally a *bla*_TEM-1B_ allele with a *P3* promoter.

Production of OXA-1 β-lactamase was identified as the likely cause of TZP resistance in seven (24.1%) of the 29 TZP-R/3GC-S isolates. Four of these isolates harboured *bla*_OXA-1_ as the only β-lactamase. Two isolates both produced OXA-1 and additionally harboured *bla*_TEM-1_ controlled by a *P3* promoter in copy numbers <2. The last OXA-1-producing isolate, WGS1364, contained both *bla*_OXA-1_ and *bla*_CTX-M-127_.

Three isolates had unknown TZP resistance mechanisms and could not be explained by our genotyping framework. Two isolates, EC180 and EC156, only harboured *bla*_TEM-135_ and *bla*_TEM-1B_, respectively, associated with *P3* promoters in copy numbers <4. Thus, no obvious cause of the observed TZP resistance was identified in these isolates. The last isolate, WGS1363, with a TZP MIC of 16, contained a previously uncharacterized CTX-M-255 β-lactamase. The sequence has been deposited at GenBank (ON876322.1).

The results above are summarized in Figure 1, in which isolates have been classified based on the most likely TZP resistance mechanism. Geometric mean of TZP MICs of grouped isolates are also indicated. The 29 isolates are presented in full detail in Supplementary Table S4.

**FIG 1.**
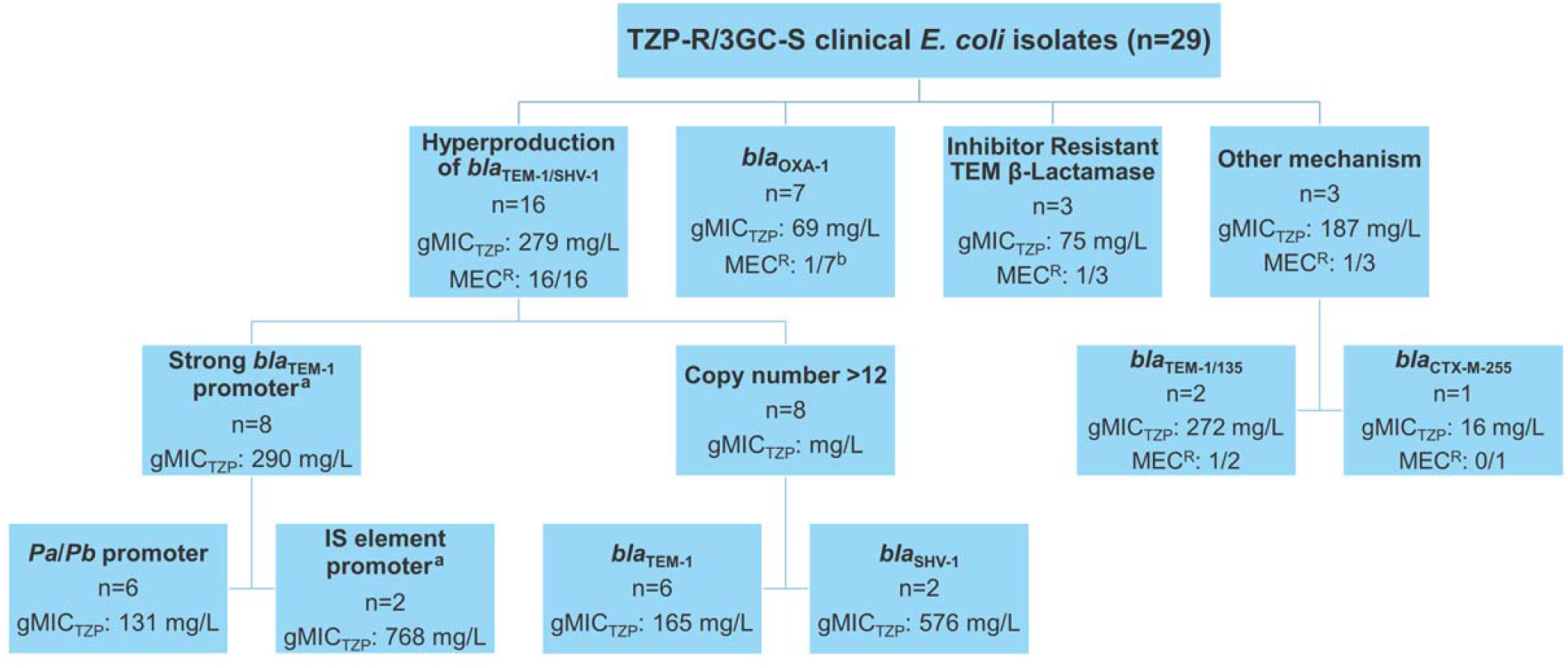
Isolates grouped by putative mechanism of TZP resistance. In three isolates, no genetic evidence for hyperproduction of TEM-1/SHV-1 or acquisition of IRT or OXA-1 β-lactamase was found. For each group containing multiple isolates the geometric mean of MIC_TZP_ (gMIC_TZP_) is indicated. MEC^R^: mecillinam resistant. ^a^Isolate EC149 both had a strong IS26/P3 hybrid promoter and a bla_TEM-1_ copy number of >12, but was assigned to the group containing strong *bla*_TEM-1_ promoters. ^b^The MEC^R^ OXA-1 isolate co-harboured *bla*_CTX-M-127_.

To further investigate the relationship between β-lactamase production and TZP-resistance, β-lactamase activity was determined in a nitrocefin conversion assay with and without prior incubation with tazobactam (Figure 2). The nitrocefin conversion activity of a TZP-susceptible isolate containing a single copy of *bla*_TEM-1_ with a *P3* promoter, EC101, was 16 mU/mg and prior incubation with tazobactam reduced this 107-fold. The 14 TEM-1 producing isolates, in which strong promoters or increased gene copy numbers were identified, had an average β-lactamase activity of 149 mU/mg (range: 37-346 mU/mg) and prior tazobactam treatment reduced this in average 46-fold. The OXA-1-producing isolates showed β-lactamase activities similar to EC101 with a range of 4-29 mU/mg and average of 15 mU/mg. However, tazobactam treatment only reduced the activities 10-fold in average. EC120 and EC126 produced both IRT and TEM-1 enzymes and showed β-lactamase activities of 106 and 231 mU/mg that were inhibited by tazobactam treatment with reduction folds of 48 and 33, respectively. EC133 produced IRT as only β-lactamase and had an intermediate β-lactamase activity of 38 mU/mg (∼2-fold the activity of EC101) that was only inhibited 6-fold by tazobactam treatment. These differences may be explained by higher activity and higher susceptibility to tazobactam inhibition of TEM-1 enzymes compared to IRTs. EC156 and EC180, in which no obvious cause of TZP resistance was identified, had high nitrocefin conversion activities (112 and 275 mU/mg, respectively) that was inhibited 88- and 93-fold by tazobactam treatment resembling isolates hyperproducing TEM-1.

**FIG 2.**
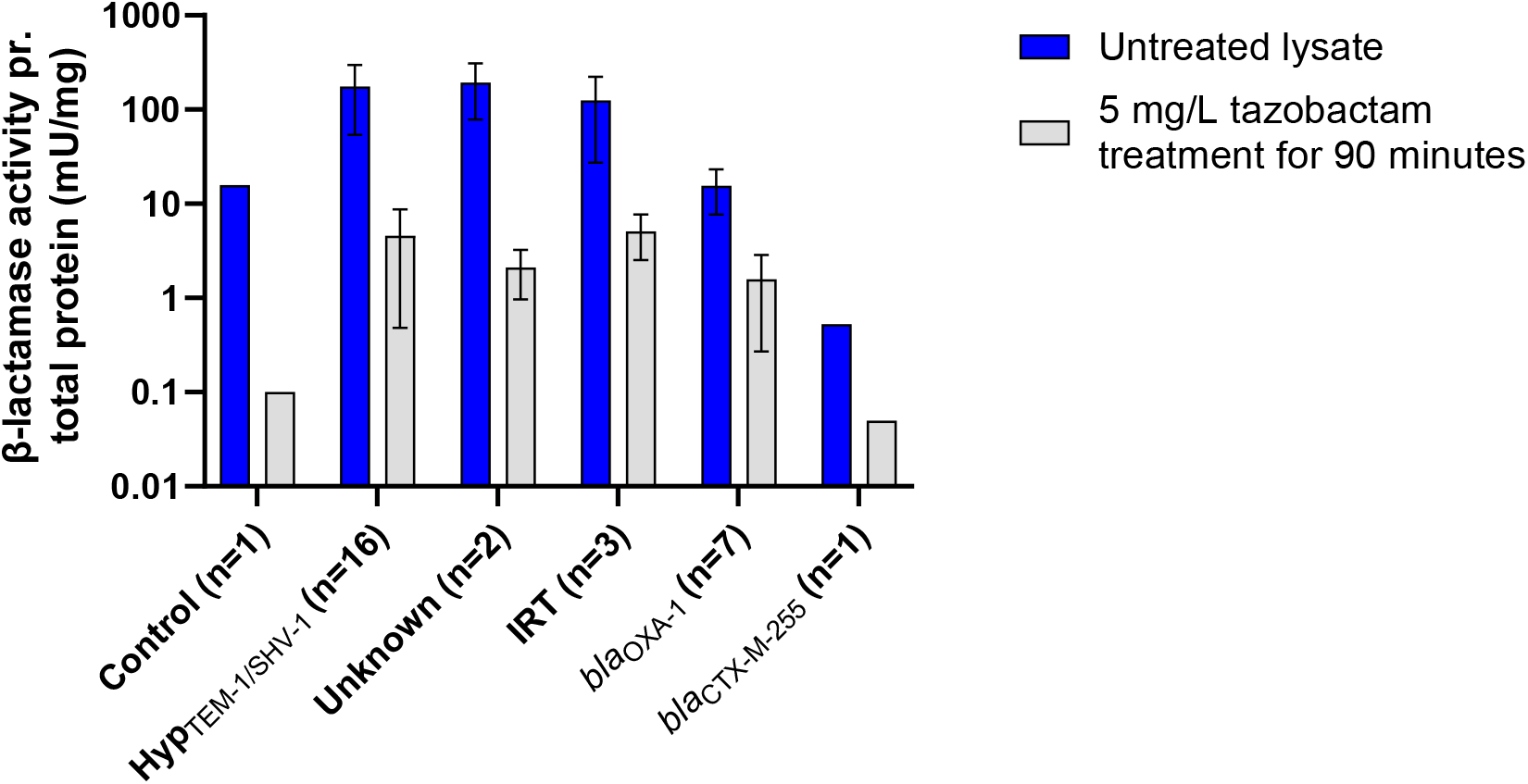
β-lactamase activity for untreated and tazobactam treated lysates of isolates grouped by different piperacillin/tazobactam (TZP) resistant mechanisms. ‘Control’ refers to the TZP susceptible bla_TEM-1_ control isolate. ‘Hyp_TEM-1/SHV-1_’ includes 16 isolates hyperproducing bla_TEM-1_ or bla_SHV-1_. ‘Unknown’ refers to the two isolates harbouring either bla_TEM-1_ or bla_TEM-135_ in low copy number and wildtype promoters but where hyperproduction is suspected. ‘IRT’ refers to isolates with an inhibitor resistant TEM β-lactamase.

WGS1363 contained a previously undescribed *bla*_CTX-M-255_ as the only β-lactamase. The nucleotide sequence of *bla*_CTX-M-255_ differs from *bla*_CTX-M-27_ by only one SNP at position 717 that translates into a G239S substitution. Hybrid assembly of Illumina and ONT sequences showed that *bla*_CTX-M-255_ was present on a 106,723 bp F1:A2:B20 plasmid. A nucleotide BLAST search of the plasmid sequence showed 98% nucleotide query coverage and >99% identity to CTX-M-27-carrying F1:A2:B20 plasmids (e.g. pH105 (CP021871) from Germany or pHP030 (LC520285) from Japan (23)). The genetic environment of *bla*_CTX-M-255_ is homologous to the environment of *bla*_CTX-M-27_. As has been reported for *bla*_CTX-M-27_ in F1:A2:B20 plasmids (24), a truncated IS*Ecp*1 element is located 42 bp upstream of *bla*_CTX-M-255_. *bla*_CTX-M-27_-containing F1:A2:B20 plasmids have been strongly associated with the *E. coli* ST131 O25:H4 *H*30R1 lineage. Consistent with this WGS1363 contains an 11,894 bp prophage, M27PP1 (LC209430), which within ST131 is specifically associated with the H30R1-M27 lineage (25).

The *bla*_CTX-M-255_-harbouring isolate had a low β-lactamase activity (0.5 mU/mg) that was reduced 11 fold upon prior treatment with tazobactam, which is comparable to that of OXA-1 enzymes and IRTs. The G239S substitution was previously obtained in a CTX-M-15 background in an error-prone PCR generated CTX-M-15 clone library with TZP selection (22). When expressed in *E. coli* TOP10 this β-lactamase conferred resistance to TZP and collateral susceptibility to cefotaxime compared to *bla*_CTX-M-15_. Therefore, we obtained the original pZS3-CTX-M-15 and pZS3-CTX-M-178 plasmids from Rosenkilde et al. and additionally cloned *bla*_CTX-M-27_ and *bla*_CTX-M-255_ with promoter in the same plasmid-backbone, pZS3, but with the native CTX-M-27 promoter to obtain the pZS3*-CTX-M-27 and pZS3*-CTX-M-255. All four plasmids together with the empty pZS3 control plasmid were expressed in *E. coli* TOP10. MICs were determined for cefotaxime and numerous penicillin/β-lactamase inhibitor combinations (Table 1). Expression of TOP10/pZS3*-CTX-M-255 reproduced the susceptibility phenotype of the parental isolate WGS1363. Both showed resistance to penicillins alone and in combination with β-lactamase inhibitors and susceptibility to cefotaxime and meropenem. Comparison of the four TOP10/pZS3(*) transformants producing either CTX-M-15, CTX-M-178, CTX-M-27 or CTX-M-255 showed that MICs of CTX-M-15 and CTX-M-27 were reduced to a larger extend by the β-lactamase inhibitors than MICs of CTX-M-178 and CTX-M-255. The ampicillin MICs were reduced from >256 to 1 or 4 mg/L for CTX-M-15 or CTX-M-27 in the presence of avibactam, whereas CTX-M-178 and CTX-M-255 were only reduced 8-fold when avibactam was added (from 16 to 2 mg/L for CTX-M-178 and from 64 to 8 mg/L for CTX-M-255. For amoxicillin, CTX-M-178 and CTX-M-255 MICs were reduced 16-fold in the presence of avibactam contra amoxicillin alone. For CTX-M-178 the piperacillin MIC was reduced from 64 to 2 (32-fold) in the presence of avibactam whereas the MIC was only reduced from >256 to 32 (>8 fold) for CTX-M-255 indicating that CTX-M-255 is more inhibitor resistant than CTX-M-178. CTX-M-255 was active against piperacillin with a MIC of >256 mg/L and MICs in the presence of clavulanic acid, sulbactam, and tazobactam of 128-256 mg/L, well above EUCAST MIC breakpoint for TZP of R>8 mg/L. For comparison, CTX-M-178 had a piperacillin MIC of 64 mg/L and a piperacillin + inhibitor MIC of 32-64 mg/L.

**Table 1.** Minimal inhibitory concentrations of TOP10/pZS3(*) transformants and parental isolate, WGS1363, containing *bla*_CTX-M-255_.

|  |  | CTX-M-15 | CTX-M-178 | CTX-M-27 | CTX-M-255 | WGS1363 | TOP10/pZS3 |
| --- | --- | --- | --- | --- | --- | --- | --- |
| Ampicillin | wo/INH | >256 | 16 | >256 | 64 | 16 | 4 |
|  | w/CLA | 16 | 16 | 128 | 32 | 8 | 4 |
|  | w/SUL | 64 | 16 | 128 | 32 | 16 | 4 |
|  | w/TAZ | 8 | 8 | 64 | 32 | 8 | 4 |
|  | w/AVI | 1 | 2 | 4 | 8 | 2 | 1 |
| Amoxicillin | wo/INH | >256 | 16 | >256 | 128 | 32 | 2 |
|  | w/CLA | 8 | 16 | 32 | 64 | 16 | 2 |
|  | w/SUL | 4 | 8 | 16 | 64 | 16 | 1 |
|  | w/TAZ | 4 | 8 | 16 | 32 | 8 | 2 |
|  | w/AVI | 0.5 | 1 | 2 | 8 | 4 | 0.5 |
| Piperacillin | wo/INH | >256 | 64 | >256 | >256 | 128 | 1 |
|  | w/CLA | 1 | 64 | 8 | 256 | 64 | 1 |
|  | w/SUL | 8 | 32 | 32 | 128 | 32 | 1 |
|  | w/TAZ | 4 | 32 | 8 | 256 | 64 | 1 |
|  | w/AVI | 1 | 2 | 2 | 32 | 16 | 1 |
| Cefotaxime |  | 256 | <0.5 | >256 | <0.5 | <0.5 | <0.5 |
| Mecillinam* |  | 0.5 | 0.25 | 0.5 | 0.25 | 0.25 | 0.25 |
| Meropenem* |  | 0.032 | 0.032 | 0.032 | 0.032 | 0.032 | 0.032 |

\*For mecillinam and meropenem, MIC was determined using E-test (bioMérieux, France). CTX-M-15 and CTX-M-178 were expressed in *E. coli* TOP10/pZS3 whereas CTX-M-27 and CTX-M-255 were expressed in *E. coli* TOP10/pZS3\*. INH=inhibitor, CLA=clavulanic acid, SUL=sulbactam, TAZ=tazobactam, AVI=avibactam, w/=with, wo/=without.

To investigate how a single amino acid substitution G239S causes this inverse resistance phenotype, the protein structure of CTX-M-255 was modelled using the webserver Robetta and its ‘comparative modelling’ option using CTX-M-27 as template. CTX-M-178 was modelled using CTX-M-15 as template in the same manner. The predicted models and their templates were visualized ‘as surface’ in PyMol (Figure 3). Residue 239 is located close to the active site of the enzyme. From the models, it is apparent that the G239S substitution limits the accessibility of the active site perhaps to an even larger extent in the CTX-M-27/255 background than in the CTX-M-15 background. This conformational change was accompanied by a change in secondary structure with an elongation of two β-sheets around the active site for both CTX-M-255 and CTX-M-178 compared to their respective template structures CTX-M-27 and CTX-M-15, see Figure S2.

**FIG 3.**
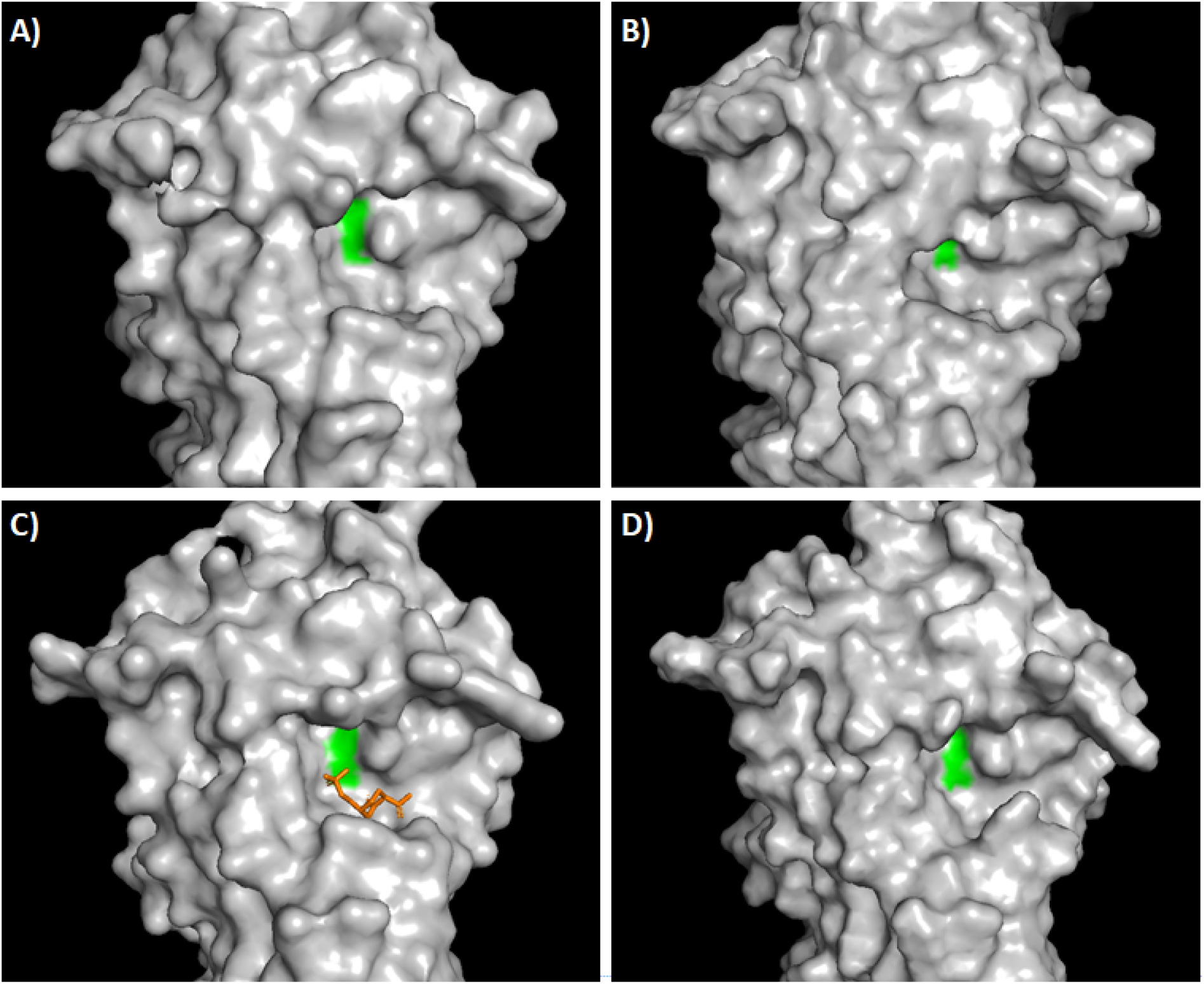
A) CTX-M-27 - PDB ID: YLP1. B) Modelled CTX-M-255. C) CTX-M-15 with avibactam bound to demonstrate the location of the active site of the enzyme. PDB ID: 4HBU. D) Modelled CTX-M-178. Position 239 is colored in green in all panels.

## Discussion

In this study, we investigated the resistance mechanisms of 29 TZP-R/3GC-S clinical *E. coli* isolates. Based on *bla* genes, their promoters and estimated copy number, we were able to identify three major resistance mechanisms as causes of TZP resistance: hyperproduction of *bla*_TEM-1/SHV_, presence of *bla*_OXA-1_ or IRT β-lactamase production, providing a genetic basis for TZP resistance in 26 of 29 isolates. By experimentally determining β-lactamase activities of the isolates, we verified hyperproduction of TEM-1/SHV-1 in 16 isolates. In addition, EC156 and EC180 appeared to hyperproduce TEM-1 or TEM-135, respectively, although the genetic basis for this was unclear. This discordance between genotype and TZP resistance phenotype was also observed in a recent study of TZP-R/3GC-S *E*.*coli* isolates that found 6.9% (n=4) of the isolates had *bla*_TEM-1_ in low copy number (<4) and associated with a weak *P3* promoter as the only β-lactamase and cause of TZP resistance identified (10). In our study, a weak *P3* promoter of *bla*_TEM-1_ but a copy number of >12 was associated with TZP resistance and 4-22 fold more β-lactamase activity than the TZP susceptible control isolate, EC101. β-lactamase activity remaining after tazobactam treatment also indicated TZP resistance in these increased *bla*_TEM-1_ copy number isolates.

We identified *bla*_OXA-1_ as the cause of TZP resistance in seven isolates (24.1%). This is similar to what was reported from Spain where the production of OXA-1 was detected as resistance mechanism in 21.0% of their amoxicillin/clavulanic acid resistant *E. coli* isolates (16). Furthermore, Edwards et al. found that 14.7% of TZP-R/3GC-S *E. coli* from a UK-wide collection carried *bla*_OXA-1_. OXA-1 isolates had measured β-lactamase activities similar to the TZP susceptible control isolate, however, after tazobactam treatment the isolates had approximately 10-fold higher β-lactamase activity than EC101. Pure OXA-1 isolates and OXA-1 with co-production of TEM-1 isolates were found resistant to TZP and amoxicillin/clavulanic acid but mecillinam susceptible. Only the isolate co-producing CTX-M-127 in addition to OXA-1 showed resistance to mecillinam which was expected since CTX-M-127 has been associated with mecillinam resistance but CTX susceptibility (22),(26). Hence, the production of OXA-1 was likely only causing TZP resistance and not mecillinam resistance consistent with the six remaining mecillinam susceptible OXA-1 isolates. IRTs was found in 10.3% (n=3) of the 29 isolates. Mecillinam susceptibility was also observed for EC120 and EC133 producing TEM-35 and TEM-30 β-lactamases. The last IRT-isolate, EC126, produced TEM-32 β-lactamase and TEM-1 from a *Pa*/*Pb bla*_TEM-1_ locus and a *P3 bla*_TEM-1_ locus was mecillinam resistant, a feature possibly attributed to concurrent TEM-1 hyperproduction. All three IRT-genes were controlled by strong promoters consistent with other findings linking IRT with strong promoters (14).

Based on this study, we propose that TZP resistant isolates hyperproducing TEM-1/SHV-1 due to a genetically detectable strong promoter or increased *bla*_TEM-1/SHV-1_ copy number relative to MLST genes are co-resistant to mecillinam. In contrast, isolates with either pure IRT β-lactamases or OXA-1 production were found mecillinam susceptible. In the clinical diagnostic laboratory mecillinam susceptibility may provide an indication of the underlying TZP resistance mechanism of TZP-R/3-GC-S *E. coli* isolates. The putative TEM-1 and TEM-135 hyperproducers EC156 and EC180 were mecillinam resistant and susceptible, respectively. The reason for mecillinam susceptibility of EC156 remains unknown.

In this study we report the phenotype associated with CTX-M-255 which when translated shares amino acid sequence with CTX-M-27 except for a G239S substitution. This β-lactamase may have previously been unrecognized since it is not associated with resistance to oxyimino-cephalosporins usually associated with CTX-M-type β-lactamases. Instead, CTX-M-255 conferred TZP resistance. MIC data of the *E. coli* TOP10/pZS3(*) transformants revealed that CTX-M-255 and CTX-M-178 with the G239S substitution are only weakly inhibited by the β-lactamase inhibitors clavulanic acid, sulbactam, tazobactam, and avibactam compared to the wildtype CTX-M-15/27 enzymes. Furthermore, CTX-M-255 and CTX-M-178 displayed low activities towards two of the tested penicillins, namely ampicillin and amoxicillin making CTX-M-255 and CTX-M-178 resembling IRT β-lactamases.

CTX-M-255 appears more active against piperacillin/avibactam than CTX-M-178, however, it is a limitation of this study that the two β-lactamases are expressed by different promoters (cf. Table S2 and Figure S1). A direct comparison is therefore not possible. Structural modelling indicates that the G239S substitution present in CTX-M-255 is associated with a narrowing of the active site cavity of the enzyme limiting access of both β-lactamase inhibitors and β-lactams like cephalosporins and to a lesser degree penicillins to the active site. As with IRT enzymes, a tradeoff between β-lactamase activity and inhibitor resistance seems to exist.

Rosenkilde et al. in an *in vitro* evolution study identified amino acid substitutions in CTX-M-15 conferring resistance to TZP and mecillinam. The substitution N135D conferred resistance to mecillinam and has been identified in clinical isolates as CTX-M-127 (https://www.ncbi.nlm.nih.gov/pathogens/isolates/#AMR_genotypes:blaCTX-M-127). The substitutions S133G and G239S in CTX-M-15 conferred resistance to TZP. These have been identified in clinical isolates as CTX-M-189 (https://www.ncbi.nlm.nih.gov/pathogens/isolates/#AMR_genotypes:blaCTX-M-189) originally reported from Illinois, USA in 2016 but now also found in UK and Canada, and CTX-M-178 originally reported from China in 2018 (KU586457), respectively. This shows that *in vitro* evolution studies can have clinical relevance. All these CTX-M-variants and the CTX-M-255 enzyme characterized here lost their hydrolytic activity to cefotaxime. In their paper, Rosenkilde et al. argued that this collateral cefotaxime susceptibility could be exploited by using combination antibiotic treatment to limit resistance evolution in clinical isolates. Clinical isolates often harbour multiple β-lactamases and sometimes multiple alleles of the same β-lactamase. In our study, 8 out of 29 isolates harboured multiple β-lactamases (cf. Table S4) and two out of three isolates carrying IRTs had maintained a TEM-1 β-lactamase. This redundancy in β-lactamase function may allow bacteria to circumvent β-lactam antibiotic combination therapy. Thus, co-expression of different β-lactamases is important in clinical isolates and could therefore be incorporated into future *in vitro* collateral sensitivity studies.

In conclusion, this TZP-R/3CG-S phenotype and the identification of another CTX-M gene that does not confer resistance to 3GC underscore that in-depth analyses of resistance mechanisms are warranted for an increasing number of isolates to improve treatment regimens and patient outcome.

